# LncRNA Bigheart trans-activates gene expression in a feed forward mechanism that facilitates calcineurin-NFAT signaling in myocardial hypertrophy

**DOI:** 10.1101/2023.12.11.571094

**Authors:** Nicolò Mangraviti, Frank Rühle, Virginie Kinet, Jana-Charlotte Hegenbarth, Ellen Dirkx, Giulia Spanò, Federica De Majo, Panagiotis Peppas, Leonie Martens, Shrey Gandhi, Robin Colpaert, Celia Rupérez, Servé Olieslagers, Inês Falcao-Pires, Martina Calore, Anika Witten, Paula da Costa Martins, Manuel Mayr, Oliver Müller, Monika Stoll, Leon J. De Windt

## Abstract

Terminally differentiated cardiomyocytes exhibit hypertrophy as a default response to injury by translating biomechanical stress into a complex network of intracellular signaling events. The molecular intricacies how calcium-dependent signaling engage molecular circuits and epigenetic modifications to activate deleterious gene programs remain enigmatic. Here we report on the re-activation of the evolutionarily conserved lncRNA “*Bigheart*”, which is repressed in the postnatal myocardium and quickly re-activated in a calcineurin-NFAT-dependent fashion in the diseased myocardium in man and mouse. In line, AAV9-mediated overexpression of lncRNA *Bigheart* in otherwise healthy primary cardiomyocytes or mouse hearts suffices to drive maladapative hypertrophy. Conversely, mice receiving a “Gapmer” antisense oligonucleotide designed to specifically silence endogenous lncRNA *Bigheart* display resistance to biomechanical stress-induced myocardial remodeling, indicating its requirement in left ventricular hypertrophy. Mechanistically, lncRNA *Bigheart* recruits the RNA binding proteins hnRNP-F1 and HMGB1 to modulate the local chromatin environment and *trans*-activate *Bigheart* target genes including *Rcan1* to stimulate calcineurin-NFAT coupling. Our observations confirm that human heart failure arises from specific susceptibilities in gene regulatory circuits that are amenable for therapeutic intervention using RNA-based therapeutics.

## INTRODUCTION

Calcineurin (protein phosphatase 2B), the calcium-calmodulin-activated, Ser/Thr protein phosphatase that dephosphorylates NFAT transcription factors,^1,2^ is essential for a wide range of biological activities, including T lymphocyte reactivity, neurological and vascular system development, bone growth, skeletal muscle fiber type switching, heart valve development, and pathological cardiac hypertrophy.^1–4^ Cardiac hypertrophy entails the maladaptive growth of heart muscle cells without an increase in cell number that occurs in response to sustained stress or injury such as pressure- or volume overload to temporally sustain cardiac output, resulting in a measurable thickening of heart muscle walls.^5^ However, cardiac hypertrophy is also accompanied by a plethora of biochemical, molecular, metabolic and extracellular changes that provoke a decrease of pump function over time rather than preserving it, predisposing the individual patient to lethal arrhythmias and overt heart failure, a serious clinical disorder that represents a primary cause of morbidity and hospitalization.^6–8^

Many signaling cascades that are necessary for myocardial hypertrophy engage molecular circuits that also control growth and gene expression in the embryonic heart, leading to the re-activation of so-called “fetal” gene programs in disease.^7^ Long ncRNAs perform a variety of regulatory roles and can recruit chromatin-modifying complexes to specific loci either near the gene encoding the lncRNA (*cis*-action), or distant target genes from the production site of the lncRNA (*trans*-action). ^9–11^ Elucidating the mechanisms how stress-responsive lncRNAs act downstream of signaling events and convert this information to epigenetic gene regulation in terminally differentiated heart muscle may unlock new convergence points for therapeutic intervention in heart failure.

## RESULTS

### *Bigheart* is an NFAT-responsive lncRNA in hypertrophic heart disease

To profile the mammalian heart for differentially expressed lncRNAs in a genome-wide fashion with lncRNA microarrays, we utilized two established mouse models with early onset cardiac hypertrophy and heart failure, one subjected to surgically induced pressure overload by transverse aortic constriction (TAC)^12,13^ and the other the calcineurin transgenic model (*Myh6*-CnA).^4^ Each model displayed substantial cardiac enlargement, fibrosis, myocyte hypertrophy and strong dysregulation of the fetal stress genes *Nppa*, *Acta1* and *Myh7* (**Fig.1a,b; Supplementary Fig.1a**). Volcano plots visualized differentially expressed lncRNAs with high statistical significance versus a robust magnitude of change in expression, yielding the surprisingly high numbers of 6,107 and 7,025 differentially expressed lncRNAs in TAC or *Myh6*-CnA hearts, respectively, compared to their wild-type controls (**Fig.1c**). A large proportion of lncRNAs (>750) were commonly down- or upregulated between both models (**Fig.1d; Supplementary Fig.1b,c**), providing more support that these lncRNAs constitute a common response of the mammalian heart to pathological growth independent of the experimental model employed.

**Figure 1.**
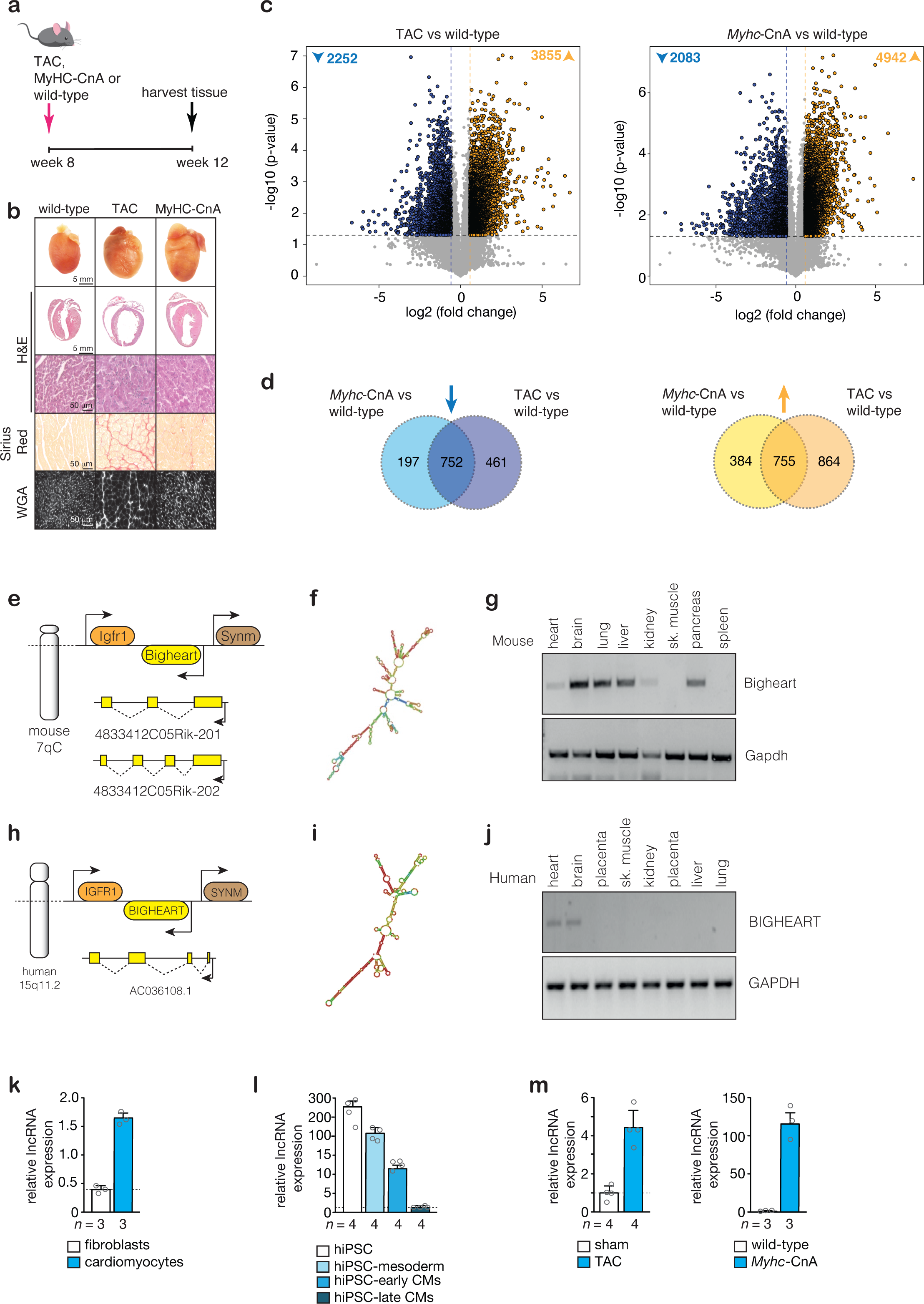
LncRNA profiling identifies “*Bigheart*” as a calcineurin/NFAT regulated non-coding RNA target gene. (a) Workflow of the experiment. **(b)** Representative images of whole hearts (top panels), haematoxylin & eosin (H&E)-stained sections of four-chamber view (second panel), high magnification H&E sections (third panel), Sirius Red stained sections (fourth panel) and wheat germ agglutinin (WGA)-stained (fifth panel) histological sections. **(c)** Volcano plots reporting the differentially expressed lncRNAs in hearts of pressure-overloaded mice compared to control animals (left panel) or in hearts of calcineurin transgenic mice compared to wild-type control animals (right panel). In blue and yellow are down- and up-regulated genes, respectively. **(d)** VENN diagrams showing the number of significantly downregulated (left panel in blue) or upregulated lncRNAs (right panel in yellow) in each heart failure model resulting from the comparison to control animals. Schematic representation of the **(e)** murine *“Bigheart”* genomic locus. **(f)** Consensus secondary structure of murine Bigheart derived from LocARNA. **(g)** RT-PCR analysis of lncRNA Bigheart expression in different murine tissues with Gapdh as loading control. **(h)** Schematic representation of the human “*BIGHEART”* genomic locus. **(i)** Consensus secondary structure of human BIGHEART derived from LocARNA. **(j)** RT-PCR analysis of lncRNA BIGHEART expression in different human tissues with GAPDH as loading control. **(k)** Real-time PCR analysis of lncRNA Bigheart in primary cardiomyocytes or fibroblasts isolated from the adult mouse heart, *n* refers to number of hearts. **(l)** Real-time PCR analysis of lncRNA Bigheart in human induced-pluripotent stem cells (hiPSCs) differentiated into mesoderm and cardiomyocytes (CMs), *n* refers to number of independent differentiations. **(m)** Real-time PCR analysis of lncRNA Bigheart in hearts from mice subjected to transverse aortic constriction (TAC) compared to sham surgery (left panel) or calcineurin transgenic mice compared to wild-type controls (right panel), *n* refers to number of hearts. Source data are provided as a Source Data file.

We uncovered a robustly dysregulated transcript in both models annotated as 4833412C05Rik-201 or ENSMUST00000181235.3 in *Mus musculus* genome assembly GRCm39, encoded on murine chromosome 7qC, and also known as “*Bigheart”* (**Fig.1e; Supplementary Fig.1b,c**). The human orthologue of the *Bigheart* lncRNA gene is encoded on human chromosome 15q11.2 and annotated as SYNM antisense RNA 1 or ENSG00000259475 in human genome assembly GRCh38. Interestingly, mouse and human lncRNA *Bigheart* show evolutionary synteny and in each case are embedded in an intragenic region between *Insulin-like growth factor 1 receptor* (*Igfr1*) and *Synemin* (*Synm*) (**Fig.1e-j**).

The conservation level on the primary structure between human lncRNA *BIGHEART* and murine *Bigheart* was relatively low (> 43%, Clustal2.1). Next, we used Minimum Free Energy (MFE) of the human and murine lncRNAs to establish secondary structure prediction using RNAfold, which revealed improved evolutionary conserved folding (**Fig.1f,i**). Although computational prediction of secondary structures is widely used, the accuracy is suboptimal and decreases for longer sequences.^14^ After successfully establishing SHAPE (selective 2-hydroxyl acylation by primer extension)- seq^15^ for the lncRNA H19 with the wildtype and the minor allele of SNP rs217727,^16^ we performed SHAPE-seq for the 588 nucleotides long sequence of human lncRNA *BIGHEART*, and obtained 3.6 to 4.4 million paired-end reads with an average coverage of 236,703, which were then incorporated during structure prediction to validate the length and secondary structure of human *BIGHEART* (**Supplementary Fig.1f,g**).

In humans, the *BIGHEART* transcript is primarily expressed in heart and brain, while in mice the expression pattern for the major isoform is more broadly present in various tissues (**Fig.1g,j**). In the heart, lncRNA *Bigheart* is more enriched in heart muscle cells rather than non-myocytes (**Fig.1k**). In human induced pluripotent stem cells (hiPSC), human *BIGHEART* is more abundantly expressed in the earlier stages of differentiation towards hiPSC-derived cardiomyocytes (hiPSC-CMs) compared to later stages of hiPSC-CM maturation (**Fig.1l**). In contrast and in line with the lncRNA profiling studies, *Bigheart* is re-expressed upon conditions of pathological hypertrophy (**Fig.1m; Supplementary Fig.1e**), reminiscent of “fetal” stress markers that are quickly re- activated in cardiac hypertrophy. Taken together, these data demonstrate that the evolutionary conserved and cardiomyocyte-enriched lncRNA *Bigheart* is higher expressed in early stages of cardiac development compared to the healthy adult myocardium and becomes strongly re-activated in response to pro-hypertrophic stimuli in the adult heart.

### LncRNA *Bigheart* is sufficient and required to induce cardiomyocyte hypertrophy

To evaluate the biological ramifications of the induction of the *Bigheart* lncRNA on the cardiomyocyte phenotype, we first considered whether the gene encoding for *Bigheart* was a direct target of the calcineurin/NFAT pathway *in vivo*, since hearts from mice with transgenic activation of calcineurin signaling displayed a very strong induction of the lncRNA transcript (**Fig.1m**). A 2 kb region upstream of *Bigheart* was scanned for evolutionary conserved cis elements representing potential NFAT-binding sites (**Supplementary Fig.2a**). Luciferase activity experiments with a *Bigheart* promoter construct harboring the NFAT consensus binding sites revealed the existence of an evolutionary conserved and functional NFAT site centered around -0.7 kb upstream of the *Bigheart* gene that was sensitive for dose-dependent titration of the VIVIT peptide^17^ (**Fig.2a,b**), providing a mechanistic basis for the observed calcineurin/NFAT responsiveness of *Bigheart* in the postnatal myocardium.

**Figure 2.**
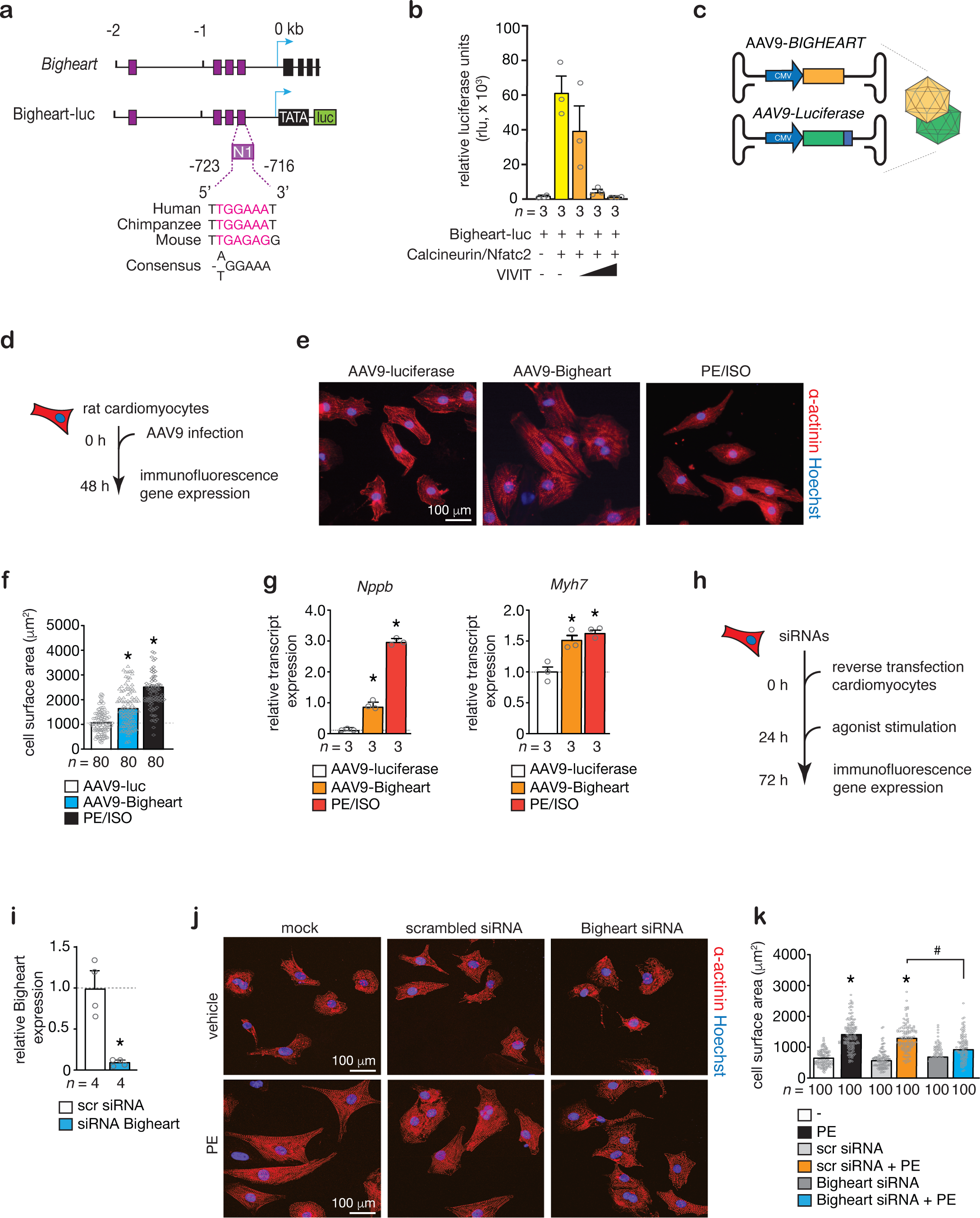
LncRNA Bigheart regulates hypertrophy in cultured primary cardiomyocytes. (a) Schematic representation of the Bigheart luciferase reporter with location of an evolutionary conserved NFAT site. **(b)** Activity assay of the Bigheart luciferase reporter construct showing the sensitivity to calcineurin/NFAT signaling, *n* refers to number of transfection experiments. **(c)** Schematic representation of serotype 9 recombinant adeno-associated vectors (AAV9). **(d)** Workflow of the experiment. **(e)** Confocal microscopy images of neonatal rat cardiomyocytes infected with AAV9- luciferase, AAV9-BIGHEART or agonist-stimulated with phenylephrine (PE) and isoproterenol (ISO); nuclei visualized with Hoechst (blue) and stained with an antibody against α-actinin (red). **(f)** Quantification of cell surface area of conditions in **(e)**, *n* refers to number of cells analyzed. **(g)** Real-time PCR analysis of fetal marker gene expression in conditions in **(e)**. **(h)** Workflow of the experiment. **(i)** Real-time PCR validation of siRNA-mediated silencing of endogenous Bigheart in primary rat cardiomyocytes, *n* refers to number of experiments. **(j)** Confocal microscopy images of neonatal rat cardiomyocytes transfected with indicated siRNAs and agonist-stimulated with phenylephrine (PE); nuclei visualized with Hoechst (blue) and stained with an antibody against α-actinin (red). **(k)** Quantification of cell surface area in conditions in (j), *n* refers to number of cells analyzed. \**P < 0.05* vs corresponding control group; #*P < 0.05* vs corresponding treatment (error bars are s.e.m.). Statistical analysis consisted of a two-tailed Student’s t-test **(i)** or One-way ANOVA followed by Dunnett multiple comparison test **(f,g,k)**. Source data are provided as a Source Data file.

Next, we made use of the high cardiac tropism and prolonged expression of serotype 9 adeno-associated viral (AAV9) vectors^18^ to evoke overexpression of the lncRNA. AAV9 vectors expressing either human lncRNA *BIGHEART*, or a control vector expressing luciferase were used to infect cultures of primary neonatal rat cardiomyocytes (**Fig.2c,d**). We performed a fluorescence microscopy-based cell surface analysis in cardiomyocytes infected with the AAV9 vectors or stimulated with the pro-hypertrophic agonists phenylephrine and isoproterenol. At 48 h, cells were stained for sarcomeric α-actinin to distinguish cardiomyocytes from non-myocytes and measured individual cell surface areas. The data show that lncRNA Bigheart overexpression suffices to evoke a cardiomyocyte hypertrophy response, albeit slightly less efficient than agonist-stimulated cardiomyocytes as measured by cell size analysis and induction of the fetal gene markers *Nppb* and *Myh7* (**Fig.2e-g**).

Conversely, we assessed the requirement of lncRNA *Bigheart* expression for agonist-stimulated cardiomyocytes to induce a full hypertrophic response. To this end, we first tested different siRNAs to provoke targeted knockdown of endogenous lncRNA *Bigheart* by RNAi *in vitro* (**Fig.2h,i**). Next, cardiomyocytes were pretreated with scrambled siRNA or siRNA against *Bigheart*. Treatment with the prohypertrophic α1- adrenergic agonist phenylephrine (PE) after scrambled siRNA transfection resulted in a robust hypertrophic response, as shown by a significant increase in cell size (**Fig. 2j,k**). RNAi to lncRNA *Bigheart* abrogated the classical hypertrophic phenotype induced by PE treatment (**Fig. 2j,k**), suggesting that *Bigheart* is required to provoke a full hypertrophic response of cardiomyocytes *in vitro*. Taken together, these data indicate that lncRNA *Bigheart* is a direct calcineurin/NFAT target gene that is both sufficient and required for cardiomyocytes to mount a full hypertrophic response.

### LncRNA *Bigheart* reactivation is required for cardiac remodeling *in vivo*

To evaluate whether a gain-of-function approach of *Bigheart* would also enhance cardiomyocyte hypertrophy *in vivo*, we injected AAV9-*BIGHEART* intraperitoneally in neonatal mice at p1 to elevate the expression and analyzed the hearts at p12 (**Fig.3a,b**). The hearts of p12 mice injected with AAV9-*BIGHEART* were significantly enlarged compared to those from mice injected with the control AAV9, increased cardiac fibrosis content and cardiomyocyte size (**Fig.3c**). Postmortem quantification further indicated that heart weight corrected for body weight and the “fetal” stress markers *Nppa*, *Nppb* and *Myh7* as hallmarks of cardiac remodeling were significantly increased in hearts of animals injected with *BIGHEART* overexpression compared to hearts of animals injected with the control AAV9-luciferase (**Fig.3d-g**).

**Figure 3.**
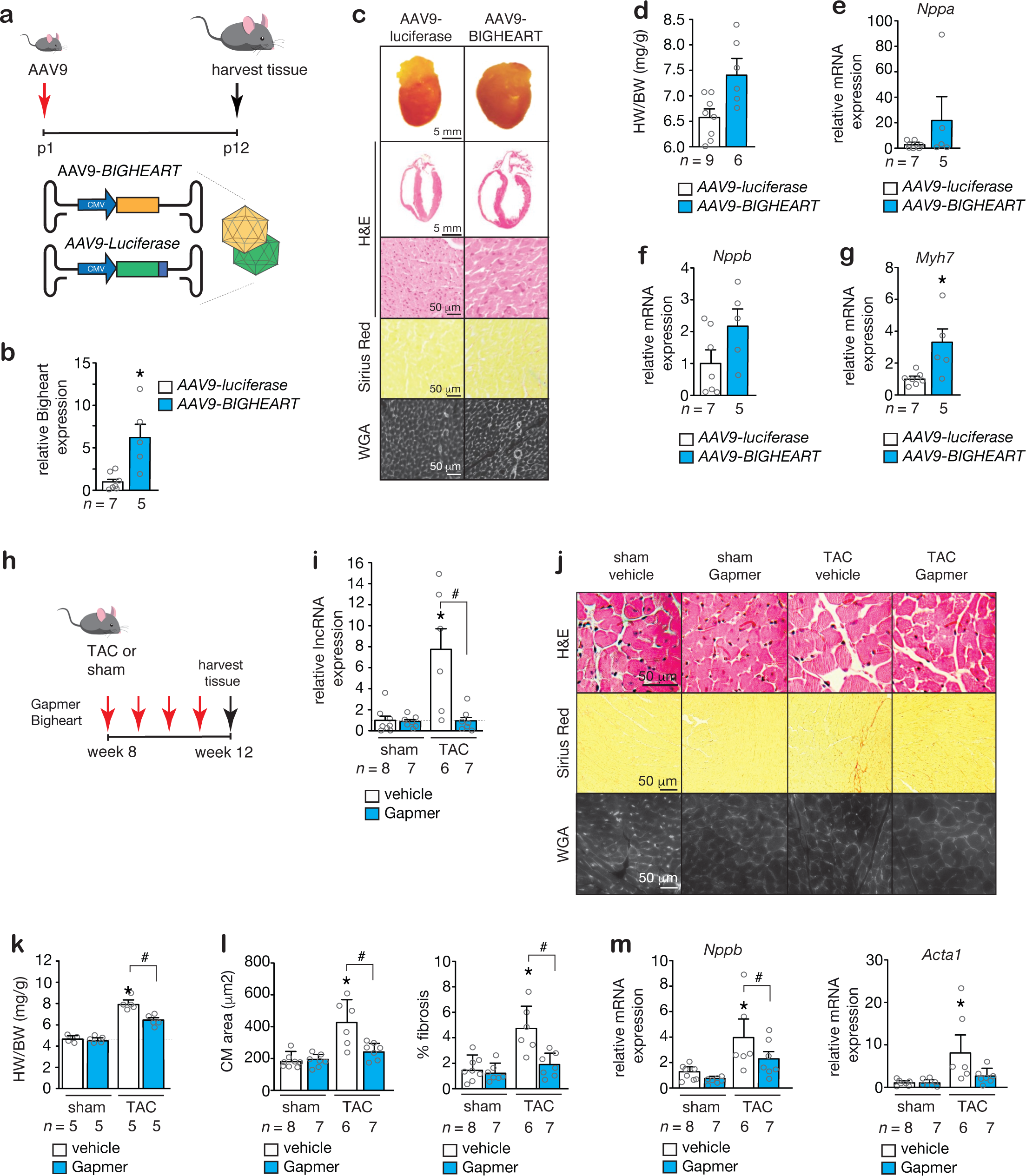
LncRNA *Bigheart* is sufficient and required to provoke cardiac hypertrophy *in vivo*. (a) Workflow of the experiment and representation of recombinant AAV9 vectors used. **(b)** Real-time PCR analysis of lncRNA *Bigheart* expression in mouse hearts infected with AAV9-luciferase or AAV9-BIGHEART. **(c)** Representative images of whole hearts (top panels), haematoxylin & eosin (H&E)-stained sections of four-chamber view (second panel), high magnification H&E sections (third panel), Sirius Red stained sections (fourth panel) and wheat germ agglutinin (WGA)-stained (fifth panel) histological sections. **(d)** Quantification of heart weight (HW)/body weight (BW) ratio, *n* refers to number of animals. Real-time PCR analysis of **(e)** *Nppa*, **(f)** *Nppb* and **(g)** *Myh7*, *n* refers to number of animals. **(h)** Workflow of the experiment. **(i)** Real-time PCR analysis of lncRNA *Bigheart* in hearts subjected to sham or TAC surgery and treated with vehicle or Gapmer. **(j)** Representative images of high magnification H&E sections (first panel), Sirius Red stained sections (second panel) and wheat germ agglutinin (WGA)-stained (third panel) histological sections. **(k)** Quantification of heart weight (HW)/body weight (BW) ratio, *n* refers to number of animals. **(l)** Histological analysis of cardiomyocyte areas and % fibrosis in hearts from (j). (m) Real-time PCR analysis of *Nppb* and *Acta1* transcript abundance in hearts subjected to sham or TAC surgery and treated with vehicle or Gapmer.\**P < 0.05* vs corresponding control group; #*P < 0.05* vs corresponding treatment (error bars are s.e.m.). Statistical analysis consisted of a two-tailed Student’s t-test **(b, d-g)** or One-way ANOVA followed by Tukey’s multiple comparison test **(i, k-m)**. Source data are provided as a Source Data file.

Finally, to assess the overall requirement of this lncRNA in experimental heart failure in the mouse, we silenced endogenous *Bigheart in vivo* with a specific Gapmer antisense oligonucleotide. We optimized dosing to weekly Gapmer administrations at a concentration of 25 mg/kg in line with previous reports to avoid toxicity effects while inducing specific and efficient loss of *Bigheart* transcripts in the heart as confirmed by real-time PCR on heart tissue (**Fig.3h,i**).^19^ We then tested the requirement of *Bigheart* in the development of cardiac disease in transverse aortic constriction (TAC) pressure overloaded hearts. To this end, wild-type mice were treated weekly with either a Gapmer antisense oligonucleotide to silence endogenous *Bigheart* or vehicle and 2 days later randomized to be subjected to TAC pressure overload for four weeks (**Fig.3h**). Cardiac size, myocyte disarray, interstitial and replacement fibrosis and cardiomyocyte size were significantly increased four weeks after TAC in vehicle-treated, control mice, but *Bigheart* silencing attenuated all these parameters of histopathological remodeling (**Fig.3j**).

Postmortem quantification further indicated that *Bigheart* silencing reduced left ventricular mass, myocyte cell size and fibrosis (**Fig.3k,l**) and attenuated re-expression of “fetal” cardiac genes *Nppb* and *Acta1* (**Fig.3m**). Taken together, these data demonstrate that lncRNA *Bigheart* is both sufficient and required for the adult heart to mount the maladaptive cardiac remodeling process following biomechanical stress.

### LncRNA *Bigheart* recruits RNA binding proteins to *trans*-activate gene expression

To understand how lncRNA *Bigheart* can evoke cardiac hypertrophy, we first performed a meta-analysis of genomic variants in the human lncRNA locus.

Cardiovascular diseases display a high proportion of genome-wide association studies (GWAS) associations in non-coding genome regions. For instance, transcript levels of the lncRNA *ANRIL* on chromosome 9p21 are directly correlated with the severity of atherosclerosis,^20^ while the lncRNA *SRA1* at chromosome 6p21 is significantly associated in dilated cardiomyopathy (DCM).^21,22^ DCM GWAS p-values were replotted for the region that surrounds human *BIGHEART* (RP11-654A16.3), but no significant association with human DCM was observed (**Supplementary Fig.3a**). A proportion of lncRNAs contain short open reading frames (sORFs) that encode for small proteins or micropeptides with largely overlooked but fundamental biological importance. We analyzed the murine and human *Bigheart* sequences with ORFfinder to verify that *Bigheart* does not code for evolutionary conserved micropeptides (data not shown).

Based upon recent classification systems, guide lncRNAs show a predominant nuclear localization and are able to recruit chromatin-modifying complexes to specific loci either near the lncRNA (*cis*-action), or distant target genes from the production site of the lncRNA (*trans*-action). We observed no induction of expression of the adjacent genes *Igfr1* and *Symn* in pressure overloaded hearts where *Bigheart* expression is strongly induced, excluding the possibility that *Bigheart* displays *cis*-action (**Supplementary Fig.3b**).

Next, to explore a possible *trans*-action for *Bigheart*, we silenced endogenous *Bigheart in vivo* with a specific Gapmer antisense oligonucleotide,^19^ induced pressure overload in mouse hearts and performed RNA-seq, which revealed a defined number of differentially expressed transcripts that were sensitive to the expression of lncRNA *Bigheart* (**Fig.4a-c**, **Supplementary Fig.3c,d**). Among the strongest induced, *Bigheart*- stimulated genes were the basic helix-loop-helix (bHLH) family member *Clock*, the z-disk located titin-assembling gene *Tcap* and the regulator of calcineurin signaling *Rcan1*.

**Figure 4.**
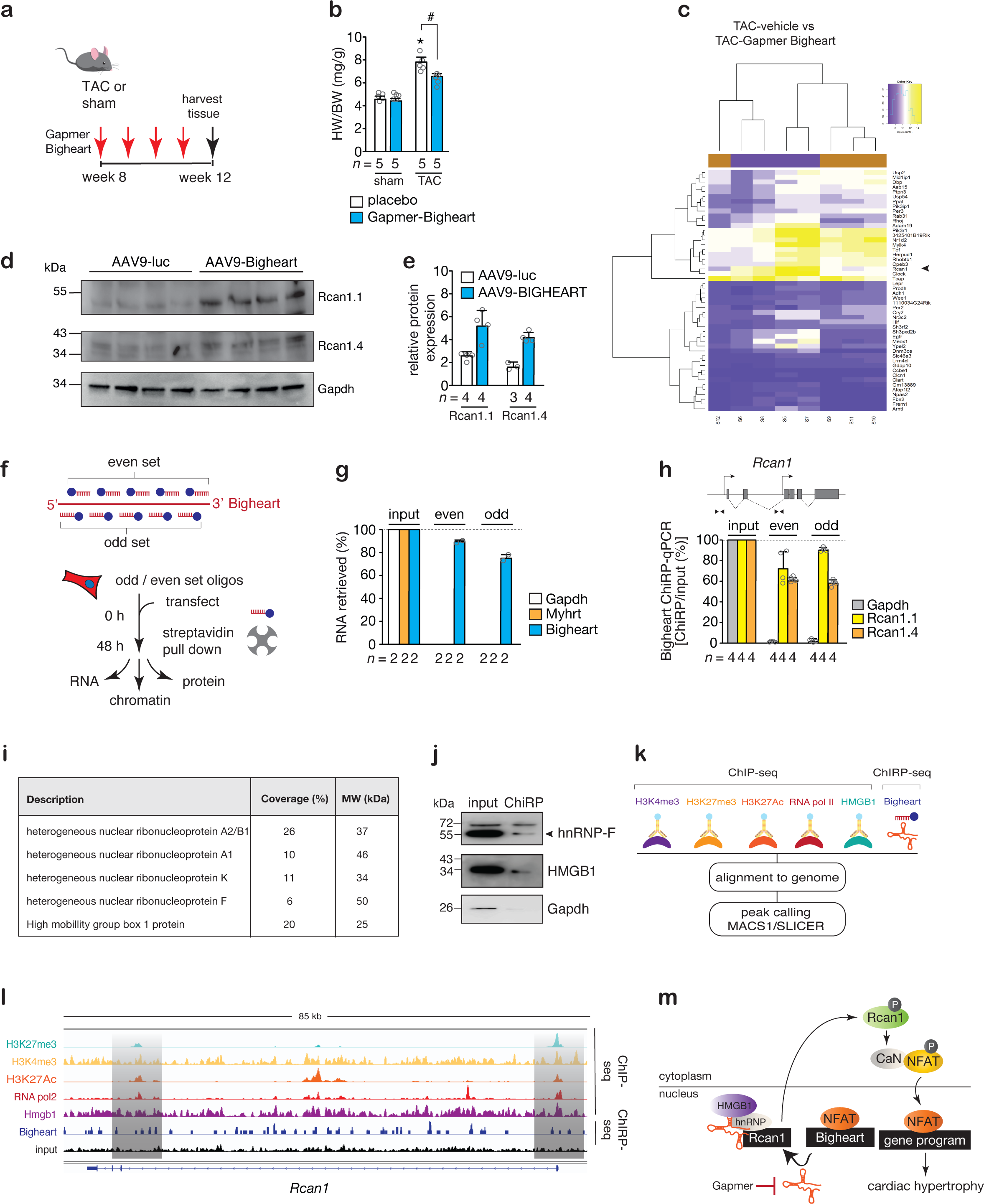
LncRNA *Bigheart* Chromatin Isolation by RNA Purification (ChIRP) reveals binding partners and chromatin targets. (a) Workflow of the study. **(b)** Quantification of heart weight (HW)/body weight (BW) ratio, *n* refers to number of animals. **(c)** Heatmap representation of myocardial transcripts differentially expressed by Gapmer-Bigheart. **(d)** Western blot analysis of endogenous Rcan1 isoforms and Gapdh as a loading control in hearts from mice receiving AAV9-luciferase or AAV9-BIGHEART, *n* refers to number of hearts. **(e)** Quantification of relative Rcan1 isoforms from (d). **(f)** Design of antisense DNA-tiling probes, grouped into ‘‘even’’ and ‘‘odd’’ sets based on their positions along mouse lncRNA Bigheart and workflow of the study. **(g)** Complementary DNA-tiling oligonucleotides efficiently and specifically retrieve lncRNA Bigheart from chromatin, *n* refers to number of independent pull-down experiments. **(h)** ChIRP-qPCR validation of peaks from Bigheart ChIRP signals in *Rcan1* promoter regions, *Gapdh* served as negative control, *n* refers to number of independent pull-down experiments. **(i)** Tabular representation of proteins detected by mass spectrometry after ChIRP retrieval of lncRNA Bigheart from chromatin. **(j)** Bigheart ChIRP retrieves hnRNP- F and Hmgb1 proteins. As a negative control, Gapdh was not detected after Bigheart ChIRP. **(k)** Worfkflow of the experiment. **(l)** ChIP-seq profiles for H3K27me3, H3K4me3, H3K27Ac, RNA Pol2, Hmgb1 and ChIRP-seq profiles for lncRNA Bigheart carried out in cardiomyocytes on the murine *Rcan1* gene. **(m)** Schematic representation of the model. \**P < 0.05* vs corresponding control group; #*P < 0.05* vs corresponding treatment (error bars are s.e.m.). Statistical analysis consisted of a One-way ANOVA followed by Tukey’s multiple comparison test **(b, e)**. Source data are provided as a Source Data file.

*Rcan1*, a member of a larger family that includes *Rcan2* and *Rcan3*,^23,24^ acts in yeast and higher organisms as an enhancer of calcineurin-NFAT signaling by virtue of its phosphorylation status and interaction with calcineurin.^25^ Given the observation that lncRNA *Bigheart* itself is also an NFAT-responsive target gene, we pursued the surprising contention that lncRNA *Bigheart* may constitute a new auto-amplification loop that stimulates calcineurin-dependent, pathological cardiac growth.

First, we could verify that both *Rcan1* isoforms increase in mouse hearts following *BIGHEART* overexpression with an AAV9 vector (**Fig.4d,e**). Since *Bigheart* works in *trans* to regulate distantly located genes such as *Rcan1*, we set out to verify the exact locations of its binding sites at higher resolution using chromatin immunoprecipitation by RNA immunoprecipitation (ChIRP)^26^ to allow high-throughput discovery of *Bigheart*-bound DNA and proteins (**Fig.4f**). In brief, mouse cardiomyocytes were transfected with biotinylated complementary oligonucleotides that tile *Bigheart*, isolated using magnetic streptavidin beads and copurified chromatin was eluted for RNA, gDNA and protein analysis. First, we verified that with both the odd- and even set of tiling oligonucleotides, we could retrieve >80% of endogenous *Bigheart* transcripts in a specific manner without contaminating unrelated RNA species (**Fig.4g**). Next, we examined whether lncRNA-associated chromatin and proteins could be copurified.

*Bigheart* ChIRP “even” and “odd” probes specifically retrieved both promoter regions of the *Rcan1* locus but not the *Gapdh* locus, demonstrating the specificity of the *Bigheart* ChIRP approach and validating *Rcan1* as a *Bigheart-*responsive target gene (**Fig.4h**). We verified by running samples on silver-stained gels and Western blotting that *Bigheart*-ChIRP efficiently purified specific proteins from mouse cardiomyocytes (**Supplementary Fig.3e,f**). Using *Bigheart* ChIRP, we interrogated by an unbiased proteomics screen what proteins are associated to *Bigheart* when bound to chromatin (**Fig.4j**). Accordingly, we detected 128 proteins in total, many of which representing various members of heterogeneous nuclear ribonucleoproteins (hnRNPs) that harbor RNA binding motifs, are predominantly expressed in the nucleus and aid in the regulation of gene expression.^27^ *Bigheart*-ChIRP also pulled down High-mobility group box 1 (Hmgb1), a non-histone DNA-binding nuclear protein and architectural chromatin-binding factor that bends the DNA and supports transcription by loosening nucleosome structure and interacting with transcription factors.^28^ By coimmunoprecipitation assays we validated that there was a direct interaction between hnRNP-F1 and Hmgb1 with *Bigheart* in mouse cardiomyocytes, but were unable to demonstrate an interaction between *Bigheart* and Gapdh, hnRNP-k, hnRNP-A2/B1 or hnRNP-C1/C2 (**Fig.4k, Supplementary Fig.3g**). Finally, to simultaneously assess lncRNA *Bigheart* occupancy and profile chromatin status in a genome-wide fashion, we performed a combined ChIRP-seq on endogenous Bigheart and ChIP-seq for H3K27me3, H3K4me3, H3K27Ac, RNA Pol2, and Hmgb1 in mouse cardiomyocytes and analyzed the murine *Rcan1* locus. Distribution analysis of H3K27Ac around the *Bigheart* peaks revealed Hmgb1 presence, chromatin activation and transcriptional activity at the two *Bigheart*-bound promoter regions of *Rcan1* in mouse cardiomyocytes (**Fig.4k,l; Supplementary Fig.3h**).

Conclusively, lncRNA *Bigheart*, which is expressed at lower levels in the myocardium during late gestation and under healthy conditions postnatally, becomes quickly re-activated upon biomechanical stress as a direct calcineurin/NFAT target gene, where it acts in *trans* by recruiting chromatin remodeling complexes to regulate distantly located genes such as *Rcan1*, thereby serving as integrative platform for auto-amplification of calcineurin-dependent signaling that drives the cardiac hypertrophy response (**Fig.4m**). Exploiting the function of this endogenous regulator of pathological growth by antisense oligonucleotide technology has therapeutic effects on cardiac remodeling in pressure overload conditions.

## DISCUSSION

Cardiomyocytes enter cell cycle arrest shortly after birth, become terminally differentiated,^29^ and exhibit hypertrophy as a default reaction to internal or external injuries by translating biomechanical stress into an intricate network of intracellular signaling events.^5–7^ Because increased ventricular mass is an independent risk factor for the development of overt heart failure and sudden cardiac death from lethal arrhythmias in humans,^30,31^ slowing hypertrophic growth may be beneficial to function or prognosis, as evidenced by studies of genetically modified mice that disrupt particular hypertrophic pathways.^12,32,33^ The stress-responsive intracellular signal transducers calcineurin and calcium/calmodulin-dependent protein kinases engage molecular circuits and epigenetic modifications such as histone or DNA modifications to re-activate gene programs that often result in cardiac remodeling as a precursor to heart failure,^34^ a serious clinical condition that is the leading cause of morbidity and hospitalization in Western societies.^6,8^

Here we describe the dynamic cardiac expression of the evolutionarily conserved lncRNA *Bigheart*, which is more abundantly expressed during embryonic development, repressed in the postnatal stage and re-activated upon calcineurin/NFAT-dependent stimulation in the human and murine adult diseased heart. Accordingly, AAV9-mediated Bigheart overexpression was sufficient to induce a maladapative hypertrophic response upon agonist stimulation *in vitro* or pressure overload *in vivo*. On the other hand, treatment with “Gapmer” antisense oligonucleotide specific for Bigheart conferred resistance to biomechanical stress-induced myocardial remodeling *in vivo*, indicating the requirement of this lncRNA in the process.

By combining a wide range of sequencing approaches and biochemical techniques, we elucidated the structural characteristics, interacting partners and molecular function of LncRNA *Bigheart* in heart muscle cells. First, computational structure prediction after selective 2-hydroxyl acylation by primer extension (SHAPE)- sequencing allowed both an experimental validation of the primary sequence as well as the secondary folding structure of this heart muscle-enriched LncRNA. Next, we could not find statistically significant associations between the presence of possible genomic variants in the *Bigheart* locus from previously executed genome-wide association studies (GWAS) in patients with dilated cardiomyopathy, suggesting that the genomic response rather than genetic variants in LncRNA *Bigheart* are likely underlying hypertrophic heart diseases. We used bulk RNA-seq in combination with Gapmer-based antisense oligonucleotide silencing of the LncRNA in the murine heart in vivo and mapped the downstream genes such as *Rcan1* that are uniquely sensitive to fluctuating expression levels of *Bigheart*. Finally, we used chromatin immunoprecipitation by RNA immunoprecipitation (ChIRP) followed by unbiased mass spec proteomics to reveal heterogeneous nuclear ribonucleoproteins (hnRNPs) and High-mobility group box 1 (Hmgb1) as preferential interacting binding partners that aid in remodeling the chromatin structure and *trans*-activation of Bigheart target genes. The functional interaction between Bigheart, Hmgb1 and hnRNPs caused an active and open chromatin structure at the two *Rcan1* promoters in heart muscle cells as experimentally revealed by a combination of genome-wide ChIRP- and chromatin immunoprecipitation (ChIP)- sequencing.

An emerging concept revolves around the ability of lncRNAs to recruit a wide variety of RNA-binding proteins (RBPs) to regulate gene expression to determine the fate of stem cells or how organs respond to internal or external cues.^35,36^ For example, hnRNPs are multifunctional RNA-binding proteins that can provoke gene expression by their interaction with lncRNAs. For example, the lncRNA *THRIL* (TNFα and hnRNP L related immunoregulatory lincRNA) complexes with hnRNP L and binds the TNFα promoter following innate activation of human macrophages,^37^ whereas the lincRNA-p21 physically interacts with hnRNP K as a coactivator for p53-dependent p21 transcription in mouse embryonic fibroblasts and various tumors.^38^ Likewise, brown fat lncRNA-1 (Blnc1) and brown adipose tissue enriched long non-coding RNA 1 (lnc-BATE1) engage in an interaction with hnRNP U to stimulate thermogenic genes and adipogenesis in brown adipocytes.^39^ Hmgb1 is an architectural non-histone chromatin-binding protein regulating transcription, DNA replication and repair, and nucleosome structure and number.^40^ Hmgb1 elicits seemingly dichotomous effects on the heart depending on its origin, subcellular localization and relative redox state, where it can mediate pressure overload-induced hypertrophy and simultaneously protect the heart muscle from excessive DNA damage caused by hypertrophic conditions.^41,42^ Interestingly, an emerging role for Hmgb1 as RNA binding protein is being recognized where in the cortex and cerebellum, Hmgb1 can associate with *brain specific DNA damage related lncRNA1* (*BS-DRL1*), where the interaction guides it to sites of DNA damage to facilitate repair in the control of motor function and purkinje cell degeneration,^43^ whereas in multiple myeloma the interaction of Hmgb1 with LncRNA *MALAT-1* controls its degradation rate and the extent of autophagic flux and tumor remission.^44^

The combined findings in this study suggest a model whereby lncRNA *Bigheart* recruits two RBPs, hnRNP-F1 and Hmgb1, to modulate the local chromatin environment and higher order chromosomal organization with simultaneously increased tri- methylation at lysine 4 and acetylation of lysine 27 in histone H3 as active enhancer marks in *Bigheart* target genes including *Rcan1* in control of the hypertrophic gene program. Given that targeted deletion of the *Rcan1/2* genes results in a loss of calcineurin signaling strength in vivo,^45,46^ the overall net function of Rcan proteins is to facilitate NFAT activity downstream of calcineurin through enhanced calcineurin-NFAT coupling in response to their increased expression as well as their posttranscriptional phosphorylation status by TGFβ-induced TAK1–TAB1–TAB2 signaling.^25^ Here, we demonstrate that re-activation of the NFAT target lncRNA *Bigheart* acts, in fact, as an RNA-based auto-amplification loop that enhances calcineurin-NFAT signaling in heart muscle cells. Previously, we identified a separate posttranscriptional feed forward mechanism whereby a calcineurin/NFAT-responsive small non-coding microRNA gene,

*miR-199b*, directly targets the nuclear NFAT kinase dual-specificity tyrosine-(Y)- phosphorylation regulated kinase 1a (Dyrk1a), effectively reducing the rephosphorylation (inactivation) of NFAT transcription factors, leading to acceleration of calcineurin-NFAT signaling strength and hypertrophic gene expression.^47^ A concept emerges where the calcineurin signaling cascade integrates with a variety of small and long ncRNA species to autoregulate kinases and accessory proteins that regulate its signaling strength in the postnatal mammalian myocardium with impact on hypertrophic gene programs. Our findings also show that human heart failure is caused by unique susceptibilities in genetic regulatory circuits, and that a better knowledge and exploitation of these circuits will aid in the prediction of future therapeutic intervention sites for RNA-based treatments.

## METHODS

### Mouse models

We used 3-6 month old calcineurin transgenic male mice in a B6SV129F1 background, which expressing an activated mutant of calcineurin in the postnatal heart under control of the 5.5 kb murine *Myh6* promoter (*Myh6*-CnA).^4^ Other mice used in this study were male and female B6SV129F1 wild-type mice of postnatal (p) day 0 or of 3-6 months of age, as well as C57Bl6/N male mice of 7-10 weeks of age (Charles River Laboratories). All animal studies were performed in accordance with local institutional guidelines and regulations and were approved by the animal review committee of Medanex Inc. and the University of Porto (0421/000/000/2018). Sample size was determined by a power calculation based upon an echocardiographic effect size. Randomization of subjects to experimental groups was based on a single sequence of random assignments. Animal caretakers blinded investigators to group allocation during the experiment and/or when assessing the outcome.

### Production of recombinant AAV vectors

Human bigheart was synthesized as MiniGene™ Synthetic Gene in pUC IDT plasmid by Integrated DNA Technologies Inc., (Leuven, Belgium) using the human reference sequence of human GRCh37 assembly. The full cDNA sequence of the lncRNA was then amplified with forward primer: 5’- GTATCATAAGGATCCCTTTCCACTGCTCTGGTGAG-3’ and reverse primer: 5’- GTATCATAAGTCGACCTCACCTAGCTGTCTGTCC-3’ and cloned into pAAV-MCS (Cat#: VPK-410, Cell Biolabs Inc.) using the restriction enzymes BamH I and HindIII sites. Recombinant AAV serotype 9 vectors were produced, purified, and titrated by real-time PCR on vector genomes at the AAV Vector Unit of the German Center for Cardiovascular Research (DZHK), Partner Site Hamburg/Kiel/Lübeck, Kiel (Germany) as described previously.^48^ B6SV129F1 mice at postnatal day 0 were intraperitoneally injected with a control AAV9 vector (AAV9-luciferase) or AAV9-BIGHEART at a dose of 1 x 10^11^ viral genome particles per animal, using an insulin syringe with 30-gauge needle. 12-15 days after injection, the hearts were collected for histological analysis.

### Aortic banding, Gapmer treatment and histological analysis

Transverse aortic constriction (TAC) or sham surgery was performed in 2-6 month-old B6SV129F1 mice by subjecting the aorta to a defined 27 gauge constriction between the first and second truncus of the aortic arch as described previously.^47,49^ All animals were randomized to receive either weekly a Gapmer antisense oligonucleotide to silence endogenous Bigheart or vehicle (phosphate buffered saline, PBS). The Gapmer specific to murine *Bigheart* was purchased at Qiagen Inc (Hilden, Germany).^19^ Treatment of vehicle or Gapmer-*Bigheart* started at 2-3 days after sham or aortic banding surgery by IP injections every 7 days (0.1 ml PBS or Gapmer-*Bigheart* at 25 mg/kg body weight/day) for a period of 5 weeks over a total of 6 weeks of pressure overload, ensuring that all animals received the last dose of vehicle or Gapmer 5-7 days before euthanasia. Six-eight weeks after surgery, animals were euthanized under deep anesthesia (sevofluorane 8%) and the hearts were removed, rinsed in ice-cold PBS, atria removed, snap-frozen in liquid nitrogen and stored in liquid nitrogen until use. A subset of hearts was arrested in diastole, perfusion fixed with 4% paraformaldehyde/PBS solution, embedded in paraffin and sectioned at 4 μm. Paraffin sections were stained with hematoxylin and eosine (H&E) for routine histological analysis; Sirius Red for the detection of fibrillar collagen; and FITC-labelled rabbit polyclonal antibody against wheat-germ-agglutinin (WGA) to visualize and quantify the myocyte cross-sectional area (1:100, Sigma Aldrich T4144). Cell surface areas and fibrotic areas were determined using ImageJ imaging software (http://rsb.info.nih.gov/ij/).

### Primary cardiomyocyte cultures, cell lines and transfections

3T3-L1 fibroblasts (ATCC, cat# CL-173) and HL-1 cells (Sigma-Aldrich, cat# SCC065) were purchased from respective vendors. 3T3-L1 were cultured in DMEM supplemented with 10% FBS, 100 units/ml penicillin/streptomycin and 2 mmol/liter L-glutamine (Thermo Fischer). HL-1 cells were cultured in Claycomb Medium (Sigma-Aldrich, cat# 51800C) supplemented with 10% FBS, 100 units/ml penicillin/streptomycin, 0.1 mmol/liter of Norepinephrine and 2 mmol/liter L-glutamine (Thermo Fischer). Cardiomyocyte cultures were isolated by enzymatic dissociation of 1 day-old neonatal rat hearts and processed for immunofluorescence as described previously.^47^ Neonatal cardiomyocytes were seeded in Corning® Primaria™ 6-well plates (for microscopy) or in Corning® Primaria™ 10 cm dishes (for RNA isolation) and one day later, cardiomyocytes were transfected with inhibitors (Dicer Substrate Duplex RNAs, Integrated DNA Technologies Inc., Leuven, Belgium) of specific lncRNA transcripts at a final concentration of 10 nM using Oligofectamine (Invitrogen). Twenty-four hours after transfection, culture medium was replaced by fresh medium; cardiomyocyte hypertrophy was induced by phenylephrine (PE, 10 μM) for an additional 24 hrs as described previously.^47^ For visualization of cardiomyocyte size and sarcomeric organization, cells were washed with ice cold PBS, fixed with 4% PFA (for 10 min) and stained with an anti-sarcomeric alpha actinin antibody [EA-53] (1:500; Abcam, cat#ab9465) followed by a phalloidin Texas Red-conjugated antibody (1:800; Molecular Probes). Nuclear staining was performed with VECTASHIELD Mounting Medium containing 4’,6-diamidino-2-phenylindole (DAPI; Vector Laboratories).

### Human induced pluripotent stem cell (hiPSC) derivation, maintenance and differentiation to cardiomyocytes

Wildtype BXS0116 hiPSCs, derived from bone marrow CD34+ cells obtained from a healthy Caucasian female donor were purchased from ATCC (ATCC® ACS-1030™). Cells were maintained in Matrigel-coated plates (Corning®), passaged with Versene solution (Thermo Fisher Scientific) and cultured in StemMACS™ iPS-Brew XF medium (Miltenyi Biotec) supplemented on the first day after passaging with 5μM ROCK inhibitor (Stemolecule Y27632). Differentiation of hiPSCs to cardiomyocytes was performed by modulation of the Wnt signaling pathway. Briefly, differentiation was started on cells cultured on Matrigel-coated plates when reaching 80- 90% confluence. Cells were initially cultured for 3 days in mesodermal induction medium, consisting of RPMI 1640 medium (Thermo Fisher) supplemented with 1% 100X glutaMAX (Thermo Fisher), 1% 100X sodium pyruvate (Invitrogen), 1% 100X penicillin/streptomycin (Invitrogen), 2% 50X B-27® Serum-Free Supplement (Invitrogen), 200uM L-ascorbic acid 2 phosphate sesquimagnesium salt hydrate (Sigma), 1uM CHIR99021 (Stemgent), 5 ng/ml Recombinant human BMP4 (R&D Systems), 9 ng/ml Recombinant Human/Mouse/rat Activin A (R&D Systems) and 5 ng/ml human FGF-2 (Miltenyi Biotec). Then, cells were cultured for 7 days in cardiac differentiation medium, consisting of RPMI 1640 medium (Thermo Fisher) supplemented with 1% 100X glutaMAX (Thermo Fisher), 1% 100X sodium pyruvate (Invitrogen), 1% 100X penicillin/streptomycin (Invitrogen), 2% 50X B-27® Serum-Free Supplement (Invitrogen), 200uM L-ascorbic acid 2 phosphate sesquimagnesium salt hydrate (Sigma) and 5uM IWP-4 (Stemgent). HiPSC-CMs were finally subjected to a 4-day metabolic selection in RPMI 1640 without glucose, without HEPES (Thermo Fisher), 1% 100X penicillin/streptomycin (Invitrogen), 2.2mM 50% sodium lactate (Sigma) and 0.1 mM 2- mercaptoethanol (Invitrogen). HiPSC-CMs were cultured for up to 25 days from the beginning of the differentiation protocol.

### Microarray analysis of lncRNAs and mRNAs

Total RNA from hearts of 3-6 months old B6SV129F1 wild-type mice, 3-6 months old *Myh6*-CnA mice, and 2-6 months old B6SV129F1 wild-type mice that received TAC surgery for 2 months (n=3 in each group) was quantified by the NanoDrop ND-1000 and RNA integrity was assessed by standard denaturing agarose gel electrophoresis. For microarray analysis, the Agilent Array platform was employed. The sample preparation and microarray hybridization were performed based on the manufacturer’s standard protocols with minor modifications.

Briefly, mRNA was purified from total RNA after removal of rRNA (mRNA-ONLY™ Eukaryotic mRNA Isolation Kit, Epicentre). Then, each sample was amplified and transcribed into fluorescent cRNA along the entire length of the transcripts without 3’ bias utilizing a random priming method. The labeled cRNAs were hybridized onto the Mouse lncRNA Array v2.0 (8 x 60K, Arraystar). After having washed the slides, the arrays were scanned by the Agilent Scanner G2505C. Agilent Feature Extraction software (version 11.0.1.1) was used to analyze the acquired array images.

### Western blot analysis

Whole tissue or cell lysates were produced in RIPA buffer (Sigma-Aldrich) supplemented with PhosSTOP- and Protease inhibitor cocktail (Roche Applied Science). Samples were boiled in 1x Leammli buffer, including 2% β-mercaptoethanol, for 5 minutes at 95°C. SDS-PAGE and Western blotting were performed using the Mini-PROTEAN 3 system (Biorad). Blotted membranes were blocked in 5% BSA / TBS-Tween. Primary antibody labeling was performed overnight at 4°C in blocking buffer. The following rabbit polyclonal antibodies were used at a 1:500 dilution: anti-HNRNPF (Sigma-Aldrich, cat# HPA069667); anti-HNRNPK (Sigma-Aldrich, cat# HPA044105); anti-HNRNPA2/B1 (Sigma-Aldrich, cat# HPA001666); anti-HNRNPC (Sigma-Aldrich, HPA051075); anti-DSCR1 (Sigma-Aldrich, cat# D6694); anti-HMGB1 (Abcam, cat# ab18256). Other antibodies applied were mouse monoclonal anti-GAPDH (1:5000, Millipore, cat# MAB374 clone 6C5), mouse monoclonal anti-alpha-Tubulin (1:5000, Sigma-Aldrich, cat# T6074) rabbit polyclonal anti-Histone H3 (1:5000, Cell Signaling Technolog, cat# 9715S) and the secondary polyclonal swine anti-rabbit immunoglobulins/HRP (1:10.000, DAKO P0399) and polyclonal rabbit anti-mouse immunoglobulins/HRP (1:10.000, DAKO P0161). Secondary HRP conjugated antibodies were applied for 1 hour at room temperature. Following antibody incubation, blots were washed for 3x10 minutes in TBS-Tween. Images were generated using the Western Lighting Ultra (Perkin Elmer) chemiluminescent detection kit and the LAS-3000 documentation system (FujiFilm Life Science). Stripping was performed using a buffer according to Abcam’s medium stripping formulation (1,5 % w/v Glycine, 0,1 % v/w SDS, 1,0% v/v Tween-20, set to pH 2,2). Output intensities were normalized for loading.

### RNA isolation

Total RNA was extracted from cultured cells or myocardial tissue of mice euthanized at the timepoints reported above using the Direct-zol RNA purification Kit (ZYMO Research) following the manufacturer’s protocol.

### cDNA synthesis

Reverse transcription of the total RNA was performed using M-MLV and RNAsin (Promega) following manufacturer’s protocol, complemented with both Oligo(dT)15 and random hexamer primers (IDT).

### Quantitative PCR

Transcriptional expression was assessed in a CFX optical thermal cycler (Biorad) from 2x SYBR Green master mix (Biorad) reactions, according to manufacturer’s instructions. Expression was normalized to expression levels of 5S rRNA. For mRNA-based reverse transcription, total RNA was reverse transcribed using hexameric random primers. The housekeeping gene ribosomal protein L7 (RPL7) was used for normalization. Fold changes were determined using the 2^-ΔΔCT^ method. Real-time PCR primer sequences used in the study are listed in **Supplementary Table 1**.

### Luciferase-reporter assays

A construct bearing 2 kb of the murine Bigheart promoter (pGL3-B.HTSS2KB) was subcloned as a HindIII and SacI fragment into the pGL3-Basic vector (Promega). pGL3-Intron3-DSCR1 harbors the complete (875 bp) of Rcan1 as described previously.^50^ Low-passage 3T3-L1 cells were grown in DMEM (Invitrogen) supplemented with 10% FCS and seeded (2,5 x 104) in 48-well plates. For cotransfection assays, 150 μg/well of either pGL3-B.HTSS2KB or pGL3-Intron3-DSCR1 reporter construct was transfected with 150 μg/well of pCDNA3-ΔCnA,and 300 μg/well pEF-BOS-hNFATp and/or 150 until 600 μg/well of pEGFP-VIVIT.^7,17^ pRL-TK (Promega; xx μg/well), containing the thymidine kinase promoter driving Renilla luciferase, was included to correct for transfection efficiency. The total amount of DNA per well was adjusted to 0 to 450 μg using pcDNA3 empty vector. The cells were washed 24 hours after transfection with phosphate-buffered saline and lysed with 100 μl of Reporter Lysis Buffer (Roche), and lysates were assayed for luciferase activity using Bright-Glo™ (Promega) on a 96-well using The VICTOR3™ Multilabel Plate Reader (PerkinElmer LAS Germany GmbH).

### Silver staining

Samples were lysate in RIPA Lysis and Extraction Buffer (thermos fish) and PhoSSTOP and protease inhibitor cocktail (roche). Then samples were boiled in 4x leammli buffer with 2% β-mercaptoethanol for 5 minutes at 95ᵒ C. then, the sample has been run in SDS-PAGE gel. Next, the gel was incubated in a Fixer solution (40% ethanol, 10% acetic acid, 50% H2O) for 1 hour. Sensitized with a solution of 0.02% sodium thiosulfate for 1 minute and incubated for 30 min with a Staining solution (0.1% AgNO3,H2O, 0.02% formaldehyde). Finally, the gel was developed in a solution of 3% sodium Carbonate and stabilized in a solution of 5% acetic acid.

### Chromatin isolation by RNA purification (ChIRP)

ChiRP was performed as described previously with minor modifications.^26^ Briefly, a series of 10 antisense DNA probes with 3’-Biotin-TEG modification were designed against the murine Bigheart full-length sequence (**Supplementary Table 2**) using LGC Biosearch Technologies software at https://www.biosearchtech.com/support/tools/design-software. 50 million HL-1 cells were cultured in Corning® Primaria™ 10-cm dishes and cross-linked in 1% glutaraldehyde for 30 min, followed by 0.125 M glycine quenching for 5 min. Next, the pellets were decanted with subsequent centrifugation and washing steps in PBS, pellets were resuspended in lysis buffer with fresh PMFS and PhosSTOP and protease inhibitor cocktail (Roche), kept on ice for 10 minutes and sonicated until the gDNA was in the size range of 100–500 bp. At this point, chromatin was diluted in hybridization buffer (500mM NaCl, 1%SDS, 100mM Tris 7.0, 10mM EDTA, 15% Formamide, add DTT, PMSF, P.I, and Superase-in fresh).), and probes were added at a final concentration of 100 pmol per 100 µL of chromatin for 24 hrs. Then 100 µL of streptavidin magnetic C1 beads per 100 pmol of probes were added and allowed to hybridize for 1 hr. Beads were then divided from the solution with a magnetic stand and washed 5 times with 1 ml wash buffer (2x SSC, 0.5% SDS, add DTT and PMSF) and processed for elution of RNA, DNA, or proteins. For the RNA extraction, beads were resuspended in 50 µL 10× RNA elution buffer (Tris 7.0, 1% SDS) and boiled for 15 min, and RNA extracted with the Direct-zol RNA purification Kit (ZYMO Research) and RNA quality assessed with controlled with nanodrop. For DNA elution, the beads were resuspended in 150 µL 3× volume of DNA elution buffer (50 mM NaHCO3, 1% SDS, 200 mM NaCl), and DNA eluted with a cocktail of 100 ug/ml RNase A and 0.1 units/microliter RNase H and samples purified with The Quick-DNA™ Miniprep Plus Kit (Qiagen). For protein elution, beads were resuspended in 3x original volume of DNase buffer (100 mM NaCl and 0.1% NP-40), and protein was eluted with a cocktail of 100 ug/ml RNase A (Sigma-Aldrich) and 0.1 units/ µL RNase H (Epicenter), and 100 U/ml DNase I (Invitrogen) at 37 °C for 30 min.

### Proteomics analysis

ChiRP proteins eluate were kept -20 °C. The gel bands were subjected to in-gel digestion with trypsin and tryptic peptides were separated on a nanoflow LC system (Dionex UltiMate 3000), eluted with a 40-min gradient, and the column (Dionex PepMap C18) coupled to a nanospray source (Picoview). Spectra were collected from an ion trap mass analyzer (LTQ-Orbitrap XL). MS/MS was performed on the top six ions in each MS scan using the data-dependent acquisition mode with dynamic exclusion enabled. MS spectra were separately analyzed in MaxQuant Software. To construct a MS/MS peak list file, up to top eight peaks per 100 Da window were extracted and submitted to search against a concatenated forward and reverse version of the UniProtKB/Swiss-Prot mouse database. A principal component analysis was performed using the normalised high/low ratios of all proteins from all samples.

Significant differences were identified using the BioConductor package Limma^51^ and Bayesian statistics to moderate variance across proteins and calculate a p-value.

### RNA sequencing

All hearts from vehicle- or Gapmer-Bigheart treated mice were prepared simultaneously during all steps of this analysis in order to exclude introduction of technical variability. Quality control of total RNA was performed using the RNA 6000 Pico Kit (Agilent Bioanalyzer) yielding in RIN values of 6.3 and higher. Removal of rRNA was carried out by use of the NEBNext rRNA Depletion Kit Human/Mouse/Rat (NEB) followed by strand-specific cDNA NGS library preparation (NEBNext Ultra II Directional RNA Library Prep Kit for Illumina, NEB). The size of the resulting library was controlled by use of a D1000 ScreenTape (Agilent 2200 TapeStation) and quantified using the NEBNext Library Quant Kit for Illumina (NEB). Equimolar pooled libraries were sequenced in a paired end mode (75 cycles) on the NextSeq 500 System (Illumina) using v2 chemistry yielding in an average QScore distribution of 84% >= Q30 score.

### Secondary structure prediction and SHAPE-Seq

We used the Vienna package RNAfold^52^ was used to predict the MFE structures for both murine and human Bigheart lncRNA. Next, we performed SHAPE-Seq for human Bigheart to determine the secondary structure. The RNA transcripts were generated in vitro from a ThermoFisher Gene Synthesis Plasmid with the T7 RNA polymerase. For this the T7 promoter and a sequence specific restriction site was attached to the sequence. For practical reasons, we restricted our approach to RNAs that are no longer than 1000 nucleotides. In vitro SHAPE-Seq was performed according to the protocol by Watters and Lucks^15^ using the 1M7 reagent, which can achieve a single nucleotide resolution. The samples were later sequenced using the Illumina NextSeq500 system and v2 chemistry in paired-end mode. The resulting reads were analyzed with Spats (v1.0.2) for generating a reactivity profile. These length-normalized reactivities were then incorporated as an additional constraint during secondary structure prediction. The SHAPE-probing information guided sampling of the structural ensemble was performed through RNAStructure^53^ pipeline consisting of Rsample, stochastic and RsampleCluster.

### Bioinformatics analyses

For analysis of the microarray datasets, quantile normalization and differential expression analysis were performed using the GeneSpring GX v12.0 software package (Agilent Technologies). After quantile normalization of the raw data, lncRNAs and mRNAs that at least 6 out of 9 samples have flags in Present or Marginal (“All Targets Value”) were chosen for further data analysis. Differentially expressed lncRNAs and mRNAs with statistical significance between two groups were identified by filtering for adjusted p-value < 0.05 and fold change >=2 or <= 0.5. Pathway analysis and GO analysis were applied to determine the roles of these differentially expressed mRNAs played in these biological pathways or GO terms, using the top 100 differentially expressed genes as it is implemented in clusterProfiler package version 3.8.1.^54^ Finally, Hierarchical Clustering on the top50 differentially expressed features was performed to show the distinguishable lncRNAs and mRNAs expression pattern among samples. For RNA-seq analysis, after quality control with FastQC v0.11.2 raw sequencing reads were trimmed for Illumina adapter sequences and quality score using trimmomatic v0.36 and aligned to the mouse reference genome GRCm38 (ensemble release 90) with hisat2 v2.1.0. Overall alignment rate was observed between 70-84% for all samples. Gene assembly was done using stringtie v1.3.4d 33 with set ‘-e’ parameter to produce read count matrices for reference genes and transcripts. Differential expression analysis of read counts was done using the DESeq2 package in R programming language. Genes are considered significant if fold changes are >= 1.5 or <= 2/3 and obtained p-values are below 0.05 after adjustment for false discovery rate (fdr). Sets of significant up-regulated and down-regulated genes were analyzed for over- represented gene ontology terms (GO) and KEGG pathways as implemented in the R-package clusterProfiler applying a significance threshold of p < 0.05 after adjustment for fdr.

## Statistics and Reproducibility

Results shown for images or blots were repeated independently at least once with similar results. The results are presented as mean ± standard error of the mean (SEM). Statistical approaches for bioinformatics analyses are described above. All other statistical analyses were performed using Prism software (GraphPad Software Inc.), and consisted of One-way ANOVA followed by Dunnet’s multiple comparison test when group differences were detected at the 5% significance level, or Student’s *t*-test when comparing two experimental groups. Differences were considered significant when *P*<0.05.

## Data availability

The data that support the findings in this study are available within the article and its supplementary information files. The complete microarray analysis was deposited at the Gene Expression Omnibus (GEO) with reference GSE236514, while the 4 SHAPSEQ and 16 mouse RNAseq data sets was submitted to National Library of Medicine (NIH) as Bioproject PRJNA991071, both in MINSEQ-compliant data submission format under restricted release date and will become publicly available pending acceptance of the current manuscript. Published resources evaluated included HomoloGene NCBI database (https://www.ncbi.nlm.nih.gov/homologene) and STRING (https://string-db.org/). Source data are provided with this paper. Any remaining raw data will be available from the corresponding author upon reasonable request.

## Acknowledgements

Next generation sequencing experiments were supported by the Core Facility Genomics of the Medical Faculty, Westfälische Wilhelms-Universität Münster. J.C.H., G.S. and L.D.W. acknowledge support from Marie Sklodowska-Curie grant agreements no. 765274 (iPlacenta) and no. 813716 (TRAIN-HEART). E.D. was supported by a VENI award 916-150-16 from the Netherlands Organization for Health Research and Development (ZonMW), an EMBO Long-term Fellowship (EMBO ALTF 848-2013) and a FP7 Marie Curie Intra-European Fellowship (Project number 627539). F.D.M. and L.D.W. acknowledge support from ERA-CVD JCT2016 EXPERT. F.D.M was supported by an ARENA-PRIME fellowship. P.D.C.M. is an Established Investigator of the Dutch Heart Foundation. L.D.W., M.C. and M.S. acknowledge support from the *Dutch CardioVascular Alliance* (ARENA-PRIME, Predict2, RACE V). M.S. acknowledges funding from the DFG (RTG2220 EvoPAD) and Marie Sklodowska-Curie grant agreement no. 813716 (TRAIN-HEART). L.D.W. was further supported by grant 311549 from the European Research Council (ERC) and a VICI award 918-156-47 from the Dutch Research Council.

## Author Contributions

N.M., V.K., E.D., S.O. performed real time PCR, transfection and cell culture experiments. N.M., V.K., S.O. performed Western blots. N.M. performed luciferase assays. N.M., V.K., and E.D. performed transcriptome analysis. G.S. and F.D.M. performed hiPSC cell cultures. N.M., R.C., C.R., P.P. and I.F.P. performed surgical procedures and histology analysis in mouse models. N.M., E.D., F.R., L.M., S.G. and J.C.H. performed bioinformatics analysis. A.W. performed next generation sequencing experiments. M.M. performed proteomics studies. O.M. generated AAV vectors. M.H., R.W., L.Z., D.N. J.V.B. and M.G. provided reagents and models. N.M., E.D. M.C., P.D.C.M., M.S. and L.D.W. designed the study. N.M., M.S. and L.D.W. wrote the manuscript. M.C, P.D.C.M., M.S. and L.D.W. acquired funding for the study.

## Competing interests

P.D.C.M. and L.D.W. are co-founders and stockholders of Mirabilis Therapeutics BV. The remaining authors declare no competing interests.

## Author Information

Correspondence and requests for materials should be addressed to M.S. or L.D.W.

## Supplementary Figures

**Supplementary Figure 1.**
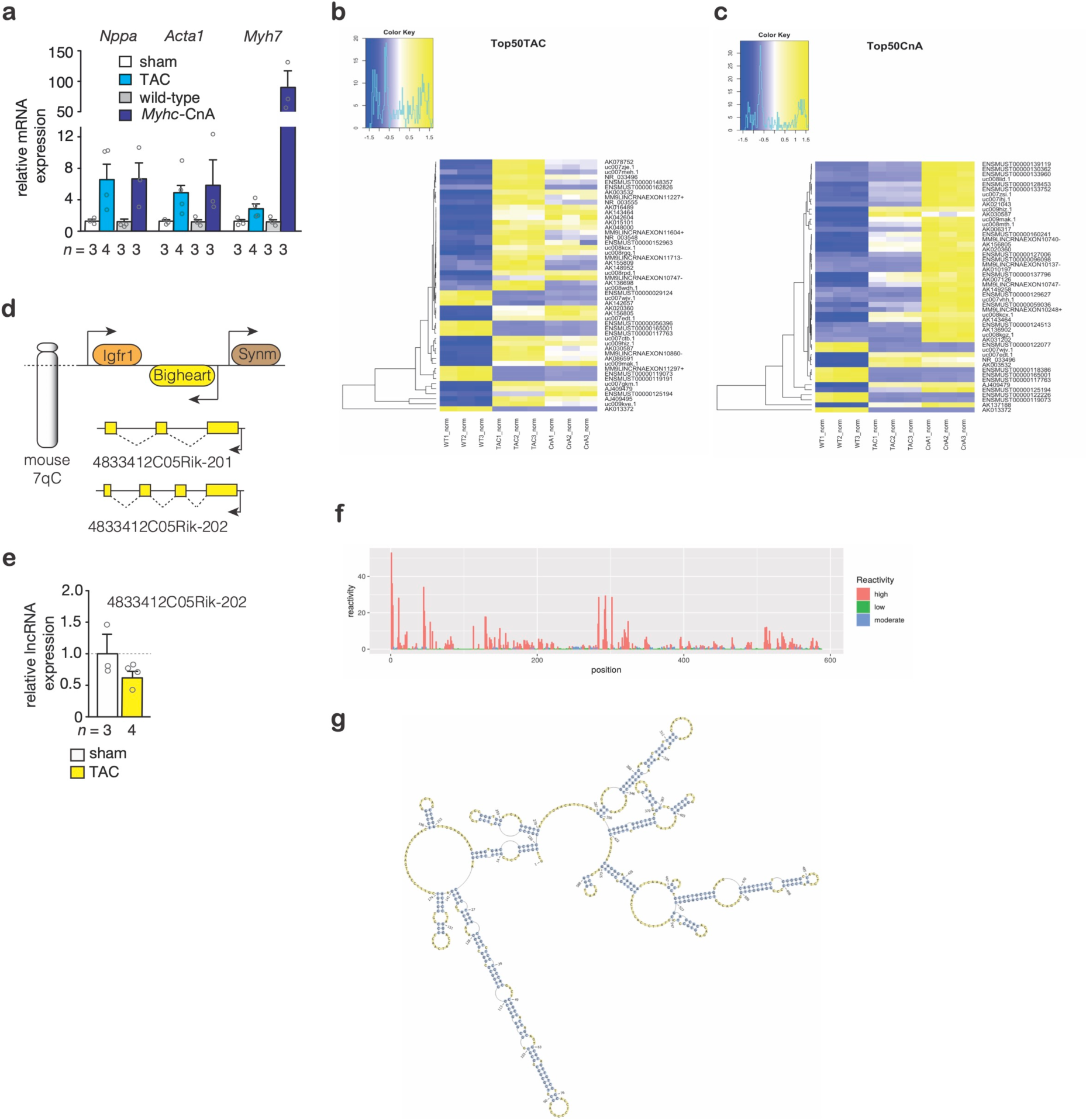
**LncRNA *Bigheart* characteristics**. **(a)** Real-time PCR analysis of fetal marker gene transcript abundance in mouse models of heart failure. **(b)** Heatmap representation of the top 50 myocardial lncRNA transcripts differentially expressed in hearts from mice subjected to sham surgery or transverse aortic constriction (TAC) for 4 weeks or **(c)** non-transgenic (nTg) or Myh6- CnA transgenic mice. (**d**) Schematic representation of the murine *Bigheart* genomic locus. **(e)** Real-time PCR analysis of the expression of its minor isoform in mice subjected to sham surgery or TAC for 4 weeks, *n* refers to the number of hearts. **(f)** Cotranscriptional SHAPE-seq reactivities from the Bigheart sequence. Color scalebars represent colorcoded ρ SHAPE-seq reactivities for each nucleotide. **(g)** Schematic of the Bigheart full-length secondary structure. Error bars are s.e.m.. Statistical analysis consisted of a two-tailed Student’s t-test. Source data are provided as a Source Data file.

**Supplementary Figure 2.**
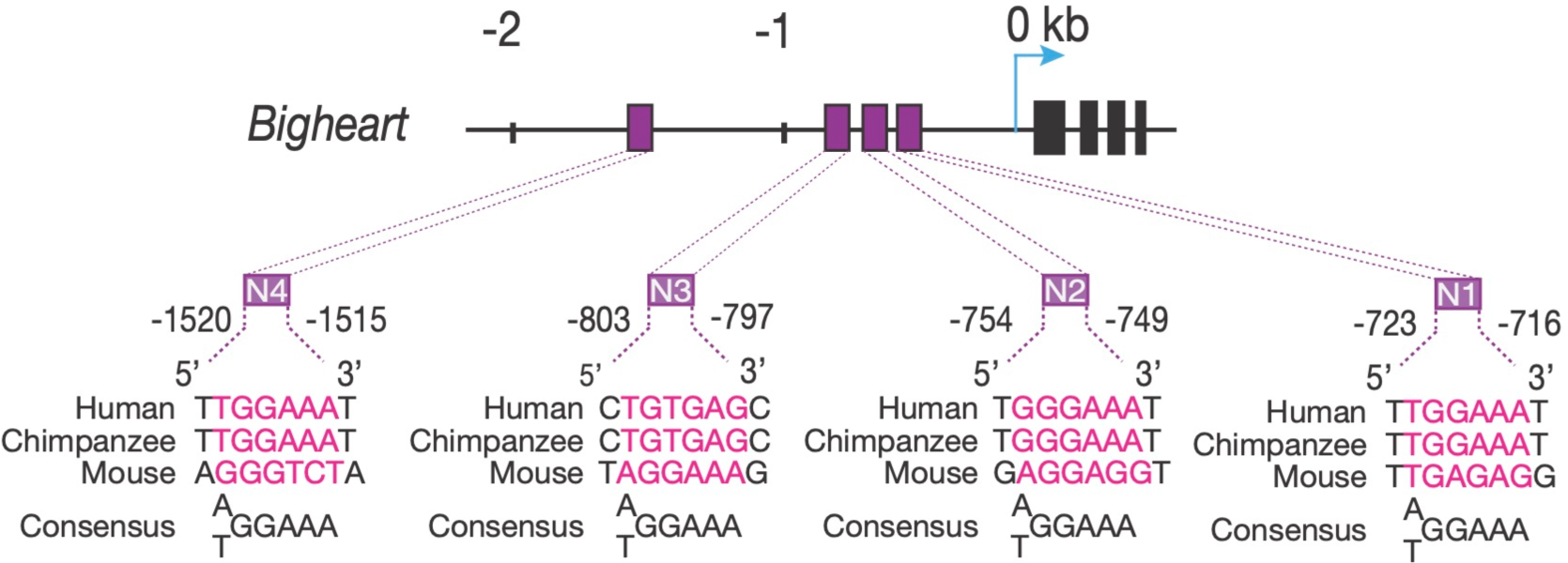
**LncRNA *Bigheart* contains several consensus binding sites for NFAT transcription factors**. Schematic representation of the murine *Bigheart* gene structure (exons represented as black squares) with the location of four evolutionary conserved NFAT sites (N1 through N4) represented as purple squares.

**Supplementary Figure 3.**
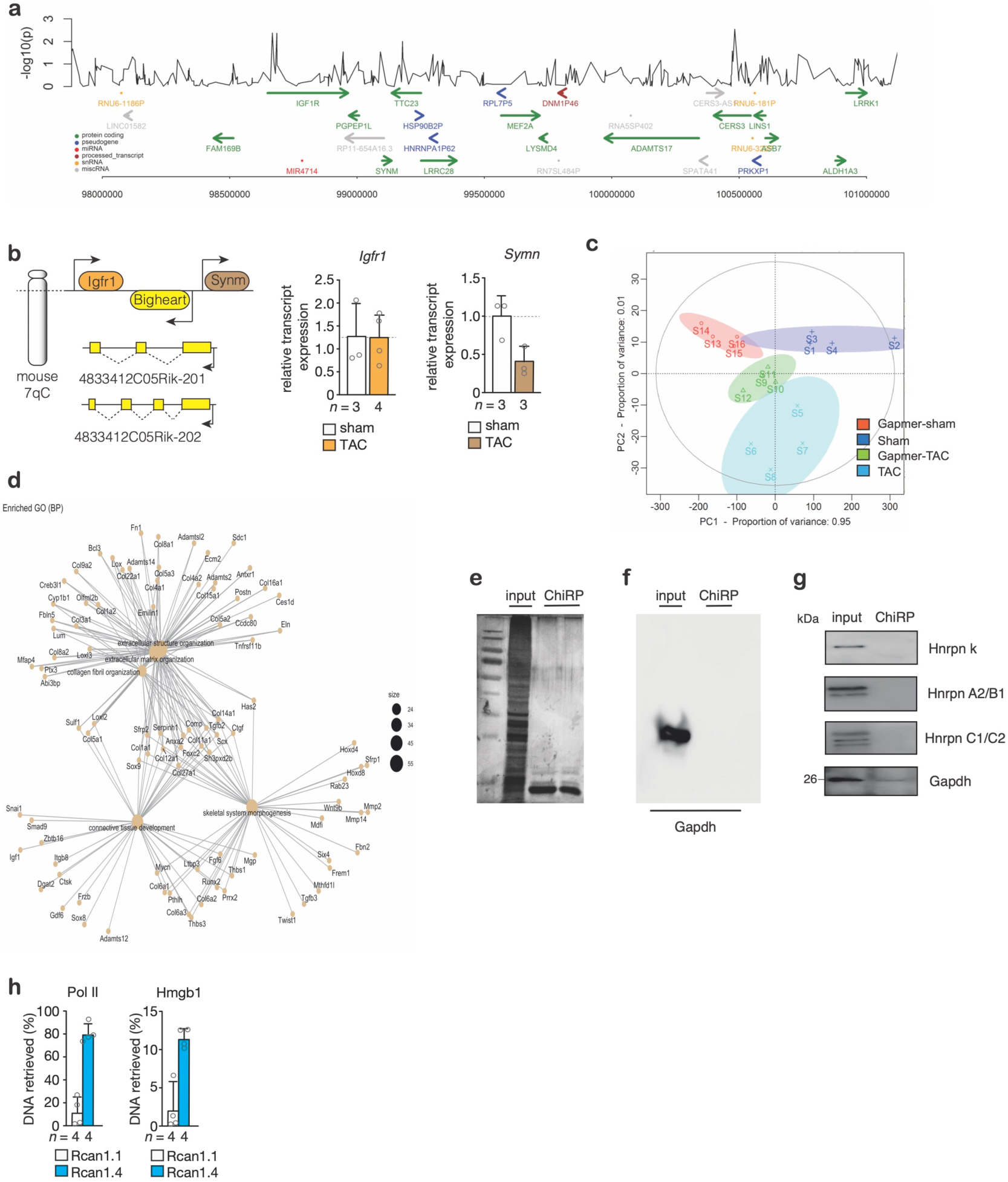
**LncRNA *Bigheart* Chromatin Isolation by RNA Purification (ChIRP)**. **(a)** Absence of statistical associations between the presence of possible genomic variants in the *Bigheart* genomic locus in genome-wide association studies (GWAS) in patients with dilated cardiomyopathy. **(b)** Schematic representation of the murine *Bigheart* genomic locus (left panel) and real-time PCR analysis of the transcript abundance of the adjacent genes *Igfr1* and *Symn* in mice subjected to sham surgery or TAC for 4 weeks, *n* refers to the number of hearts. **(c)** PCA plots of the mRNAs data representing the expression profiles trends of hearts from mice subjected to transverse aortic constriction (TAC) and treated with vehicle or Gapmer for 4 weeks. The plots have been generated using the first two principal components, which together account for the 95% and 1% of the variance between samples, n=3 hearts for each mouse model. **(d)** Gene-Concept networks visualizing the individual protein-coding genes associated to the GO-terms found enriched for Gapmer-treated mice. (e) Silver-stained PAGE gel with visualization of total proteins before or after Bigheart-ChIRP. **(f)** Bigheart ChIRP does not retrieve Gapdh or **(g)** hnRNP-k, hnRNP-A2/B1 or hnRNP-C1/C2. As a negative control, Gapdh was not detected after Bigheart ChIRP. Error bars are s.e.m. Statistical analysis consisted of a two-tailed Student’s t-test. Source data are provided as a Source Data file.

**Supplementary Table 1.**
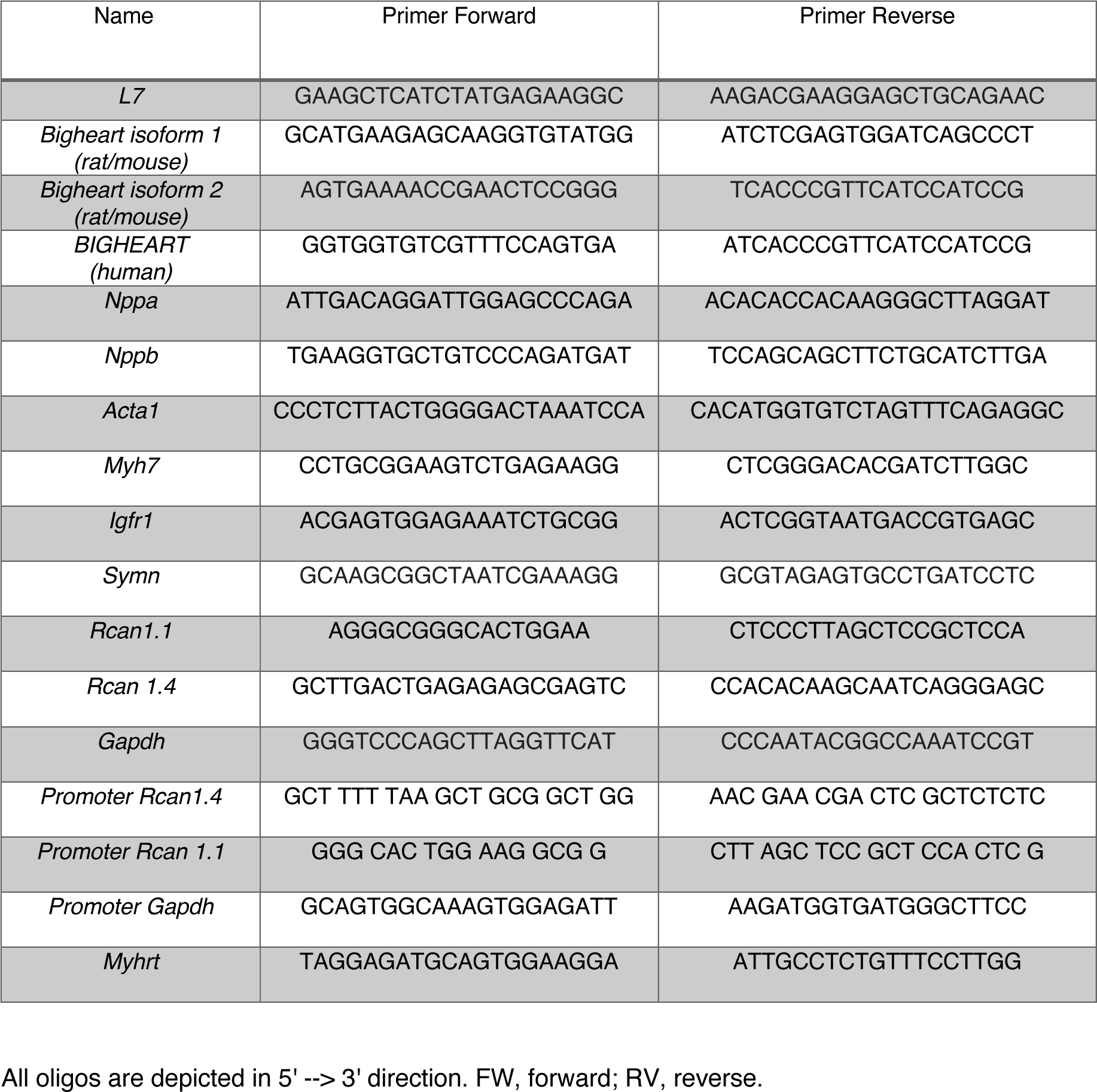
real-time PCR primers used in the study.

**Supplementary Table 2.**
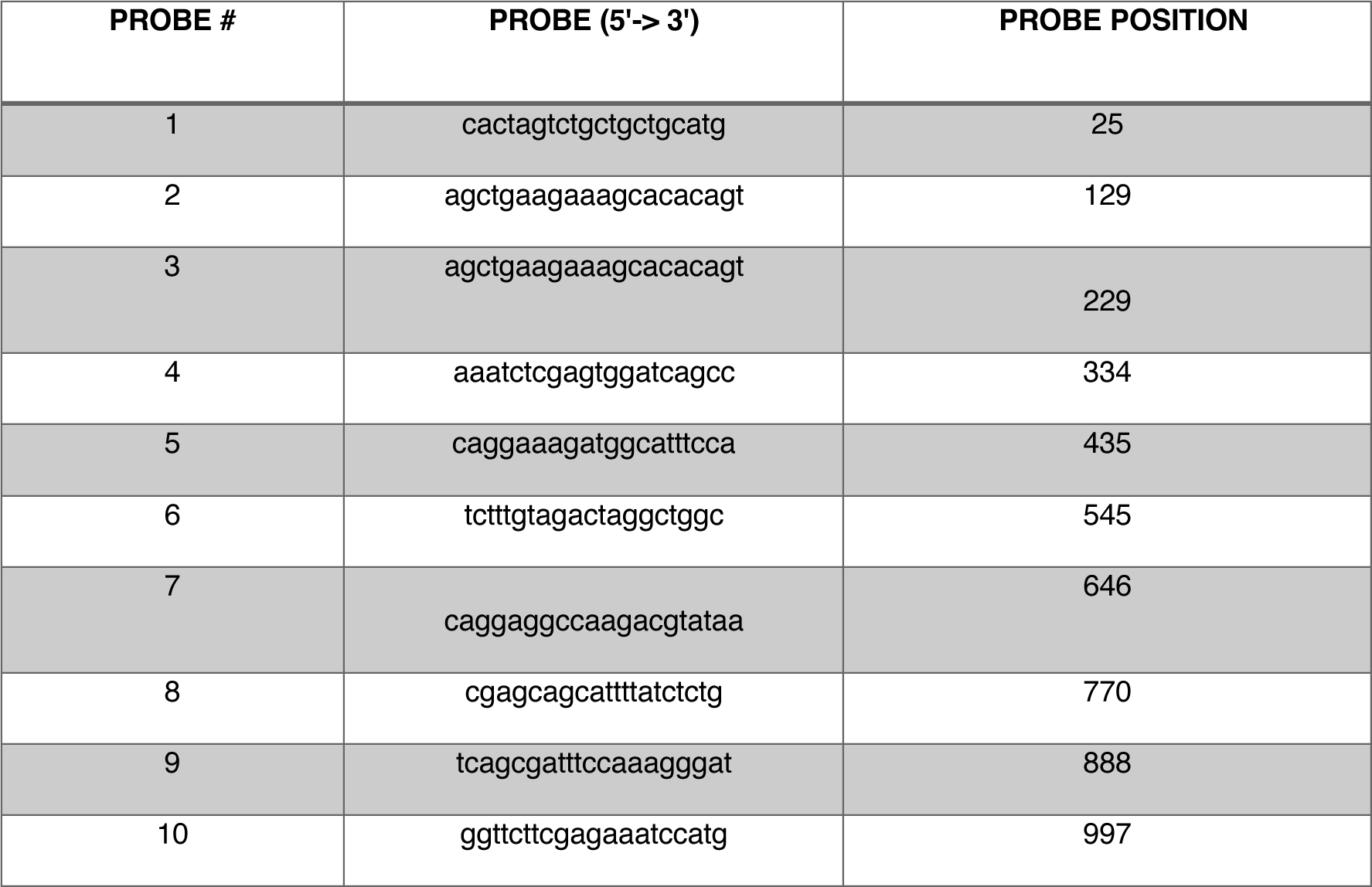
Antisense DNA-tiling probes, grouped into ‘‘even’’ and ‘‘odd’’ sets based on their positions along mouse lncRNA *Bigheart* used for Chromatin Isolation by RNA Purification (ChIRP).

